# Subthalamic nucleus local field potentials recordings reveal subtle effects of promised reward during conflict resolution in Parkinson’s disease

**DOI:** 10.1101/454173

**Authors:** Joan Duprez, Jean-François Houvenaghel, Thibaut Dondaine, Julie Péron, Claire Haegelen, Sophie Drapier, Julien Modolo, Pierre Jannin, Marc Vérin, Paul Sauleau

## Abstract

Cognitive action control depends on cortical-subcortical circuits, involving notably the subthalamic nucleus (STN), as evidenced by local field potentials recordings (LFPs) studies. The STN consistently shows an increase in theta oscillations power during conflict resolution. Some studies have shown that cognitive action control in Parkinson’s disease (PD) could be influenced by the occurrence of monetary reward. In this study, we investigated whether incentive motivation could modulate STN activity, and notably STN theta activity, during response conflict resolution. To achieve this objective, we recorded STN LFPs during a motivated Simon task in PD patients who had undergone deep brain stimulation surgery. Behavioral results revealed that promised rewards increased the difficulty in resolving conflict situations, thus replicating previous findings. Signal analyses locked on the imperative stimulus onset revealed the typical pattern of increased theta power in a conflict situation. However, this conflict-related modulation of theta power was not influenced by the size of the reward cued. We nonetheless identified a significant effect of the reward size on local functional organization (indexed by inter-trial phase clustering) of theta oscillations, with higher organization associated with high rewards while resolving conflict. When focusing on the period following the onset of the reward cue, we unveiled a stronger beta power decrease in higher reward conditions. However, these LFPs results were not correlated to behavioral results. Our study suggests that the STN is involved in how reward information can influence computations during conflict resolution. However, considering recent studies as well as the present results, we suspect that these effects are subtle.

## 1. Introduction

Cognitive action control is a subset of cognitive control involved in our daily life ability to adapt our motor behavior to a changing environment and to our intentions (Ridderinkhof et al., 2011). It is especially important when facing a situation involving multiple potential behavioral outcomes that conflict with each other. In cognitive neuroscience, this process is traditionally measured using computerized conflict tasks, such as the Simon task or the Eriksen flanker task (van den Wildenberg et al., 2010; Hommel, 2011). In these experiments, the participants have to overcome an automatically activated behavior that conflicts with the correct action. An increase in reaction time (RT) and errors has been invariably described in the case of conflict situations, and these effects have been extensively used as metrics of cognitive action control performance, both in healthy (van der Lube & Verleger, 2002; Forstmann et al., 2008; Duprez et al., 2016), and in patient populations (Georgiou-Karistianis et al., 2007; Cespon et al., 2015; Riesel et al., 2017; Duprez et al., 2017).

The cognitive action control process relies on several cortical areas, with accumulating evidence arguing for a fronto-parietal circuit of conflict-processing (Cohen & Ridderinkhof, 2013; Cohen & van Gaal, 2014; van Driel et al., 2015). The pre-SMA seems systematically involved, as shown by the consistent increase in both neuronal oscillations theta power (4-8 Hz) (see Cohen, 2014 for a review), and BOLD activity when conflict arises (Förstmann et al., 2008). Other cortical regions are also involved in cognitive action control performance, such as the right inferior frontal cortex (rIFC) or anterior cingulate (see Ridderinkhof et al., 2011 for a review). Cognitive action control also depends on subcortical structures represented by the basal ganglia (BG). Their role is assumed to be crucial, since they are known to form cortical-subcortical loops involved in motor behavior, cognition, and emotional processing (Haber, 2014; Jahanshahi et al., 2015; Péron et al., 2013; Argaud et al., 2018).

The involvement of the BG in cognitive action control has notably been inferred on the basis of observed lack of cognitive action control. This has been repeatedly described in Parkinson’s disease (PD) patients, in which one of the BG, the substantia nigra *pars compacta*, shows a gradual degeneration of dopamine neurons. This results in a dysregulation of cortical-subcortical loops, ultimately leading to disabling motor and non-motor symptoms. PD patients present impairments in performing tasks such as the Simon task, with a stronger effect of conflict on both RT and accuracy as compared to healthy participants (Wylie et al., 2009, 2010; Duprez et al., 2017). Behavioral results and computational modeling studies have led to the proposal that the STN inhibits responses during conflict situation to avoid impulsive action selection (Frank, 2006; Bogacz & Gurney, 2007; Bonnevie & Zaghloul, 2018). Although behavioral results strongly suggest the involvement of the BG, more direct evidence came from STN recordings that are possible following electrodes implantation for deep brain stimulation (DBS). Recordings can indeed be performed in the interval between surgery and the implant of the subcutaneous stimulator, thus enabling the recording of STN local field potentials (LFPs) while the patient performs a cognitive task. STN LFP studies strengthened the arguments in favor of its key implication in cognitive action control, by describing the dynamics of neural oscillations inferred through time-frequency based analyses of the LFPs. The most significant and replicated result is that conflict situations yield an increase in STN theta band power (Brittain et al., 2012; Zavala et al., 2013; Cavanagh et al., 2011). Evidence of functional connectivity between the pre-SMA and STN in the theta band, as well as evidence of correlation between power and reaction time, have led to the proposal that the medial prefrontal cortex drives STN inhibitory activity during cognitive action control (see Zavala et al., 2015 for a review).

Since the STN is thought to be a key integration structure in the BG, recent studies have not only focused on its role in cognitive action control and inhibition abilities, but also on its computational role regarding motivation and reward processing. These studies are also of critical relevance, since the dopaminergic system enabling harmonious neural communication within the BG is also a core system in reward processing (Wise, 2004). For instance, STN involvement has been shown by studies revealing that STN DBS can modulate reward processing in PD patients (Wagenbreth et al., 2015). Since most of our daily behaviors are motivated, an important aspect is the influential role of motivation, and more specifically of incentive stimuli, on cognitive action control performance. It is indeed fairly accepted that motivational stimuli can trigger reward expectation that, in turn, influences decision making (Berridge, 2004). Recent studies have accordingly investigated whether incentive stimuli could modulate cognitive action control performances, mainly relying on motivated conflict tasks in which an incentive stimulus is presented before the imperative stimulus, thereby triggering reward expectation. So far, no strong consensus has been achieved on the influence of such stimuli on the ability to resolve conflict. Indeed, some studies in healthy participants described a beneficial effect of the expected reward, and measured as a smaller effect of conflict on behavioral measures (Padmala & Pessoa, 2011); while others found a detrimental effect (Padmala & Pessoa, 2010; Houvenaghel et al., 2016a) or no significant influence (Aarts et al., 2014; van den Berg et al., 2014). Finally, a recent study has reported that the STN was involved in the influence of promised rewards on conflict resolution since patients with STN-DBS showed more impulsive responses in low rewarded conflict context compared to patients without DBS (Houvenaghel et al., 2016b).

So far, only a few studies have described the STN oscillatory activity while treating a monetary reward. For instance, Zénon et al (2016) investigated STN response to a promised monetary reward during an effortful behavior. The authors showed that STN low-frequency oscillatory power (< 10 Hz) was modulated by the size of a promised monetary reward, with increased power when the reward was high. In another study focusing on gambling in PD patients, the authors proposed a task in which patients, with or without pathological gambling, had to choose a stimuli that could represent either a monetary loss or a gain with different probabilities (Rosa et al., 2013). In that study, modulation of STN oscillatory power in response to a potential reward was also described in the low frequencies (< 12 Hz). Taken together, even if further studies are needed to confirm the STN response to a motivational stimulus, evidence is pointing at an involvement of low frequencies, notably in the theta range, as it is the case for conflict processing.

The precise mechanisms through which the STN acts in cognitive action control and in reward processing are still elusive. However, there is convincing evidence that the STN plays a role in controlled behaviors, most of which are directed to specific goals and under the influence of motivation, another process known to involve the STN (Bonnevie & Zaghloul, 2018). However, to our knowledge, no study has described the involvement of the STN in how reward stimuli modulates cognitive action control performance. In this study, we analyzed STN LFPs recorded in 16 PD patients performing a Simon task motivated by the presentation of promised monetary reward before the imperative stimulus. We hypothesized that STN oscillatory activity is modulated by the reward both at reward presentation, and during conflict processing, in the theta band. More specifically, we sought to verify that STN theta power increases according to the reward size, and that the conflict-related increase in theta power would also be modulated by the reward size.

## 2. Methods

### 2.1 Patients and surgery

Sixteen patients (9 women) with idiopathic PD, as defined by Parkinson’s UK Brain Bank (Hughes et al., 1992), took part in this study. All patients underwent bilateral STN DBS at Rennes University Hospital (France) and were selected for surgery based on standard criteria (Welter et al., 2002). STN DBS was indicated because of disabling motor symptoms occurring in spite of optimum oral therapy. Patients underwent a battery of neuropsychological tests that revealed no major attentional or executive dysfunctions. Patients’ clinical details are summarized in Table 1. Targeting and bilateral implantation of stimulation electrodes was done as previously described (Péron et al., 2017). The electrodes model used were Medtronic model 3389 (Medtronic Neurological Division, Minneapolis, MN, USA) with four platinum-iridium cylindrical surfaces (1.27 mm in diameter and 1.5 mm in length) and a contact-to-contact separation of 0.5 mm. Contact 0 was the most ventral and contact 3 the most dorsal. Lead location was confirmed by a three-dimensional CT scan acquired a few days after implantation.

**Table 1:**
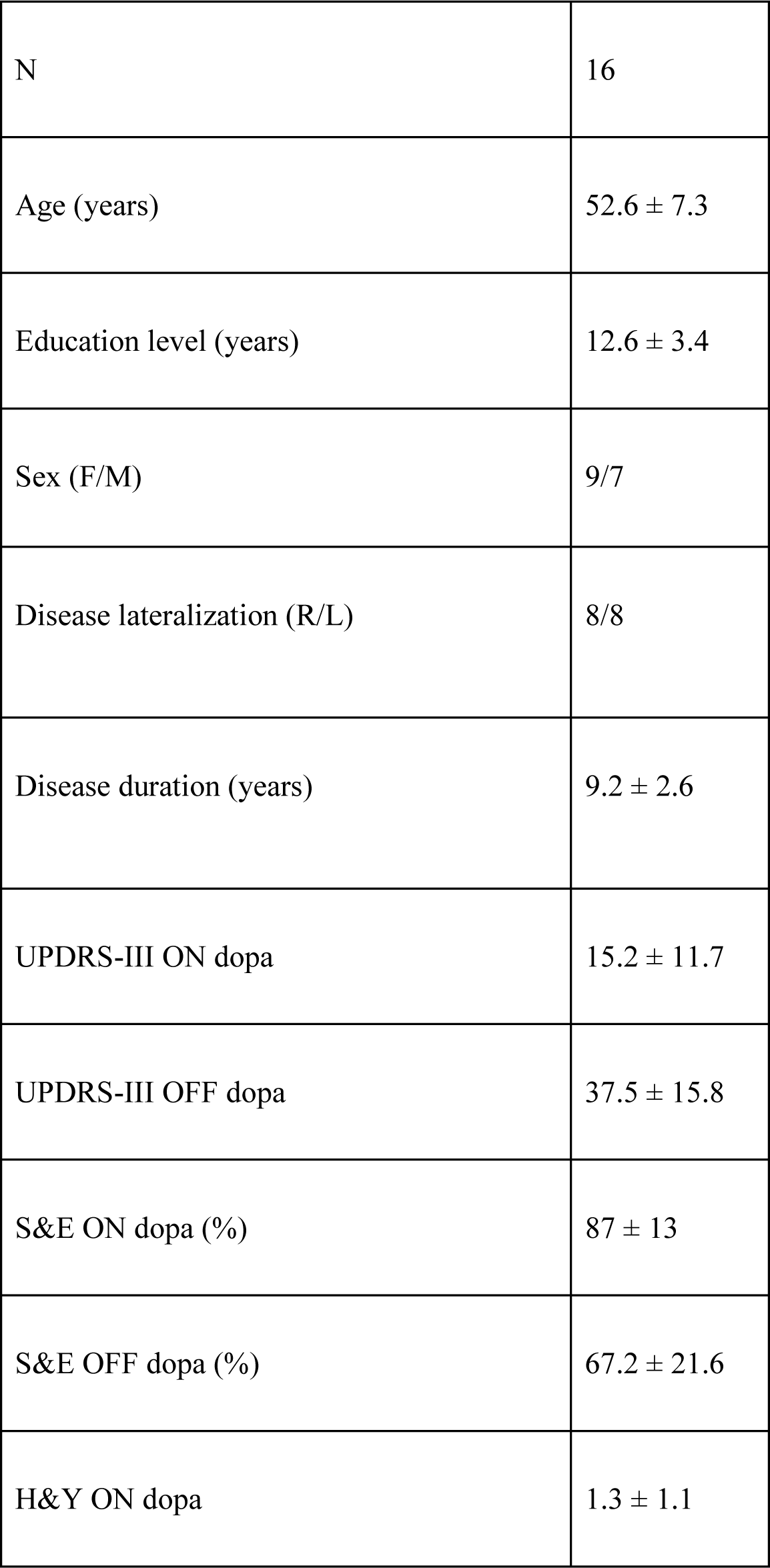
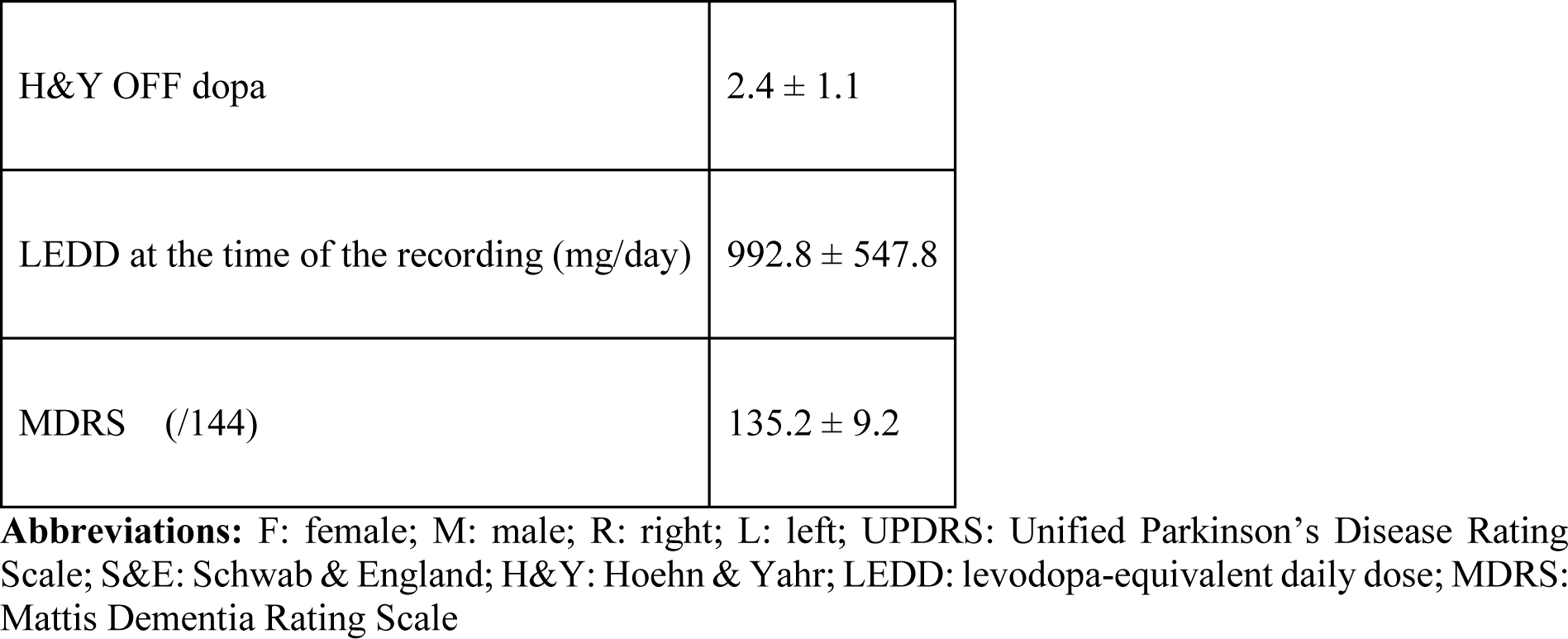
Patients’ preoperative clinical characteristics (average ± standard deviation).

|  |  |
| --- | --- |
| N | 16 |
| Age (years) | 52.6 ± 7.3 |
| Education level (years) | 12.6 ± 3.4 |
| Sex (F/M) | 9/7 |
| Disease lateralization (R/L) | 8/8 |
| Disease duration (years) | 9.2 ± 2.6 |
| UPDRS-III ON dopa | 15.2 ± 11.7 |
| UPDRS-III OFF dopa | 37.5 ± 15.8 |
| S&E ON dopa (%) | 87 ± 13 |
| S&E OFF dopa (%) | 67.2 ± 21.6 |
| H&Y ON dopa | 1.3 ± 1.1 |
| H&Y OFF dopa | $2.4 \pm 1.1$ |
| LEDD at the time of the recording (mg/day) | $992.8 \pm 547.8$ |
| MDRS (/144) | $135.2 \pm 9.2$ |

**Abbreviations:** F: female; M: male; R: right; L: left; UPDRS: Unified Parkinson's Disease Rating Scale; S&E: Schwab & England; H&Y: Hoehn & Yahr; LEDD: levodopa-equivalent daily dose; MDRS: Mattis Dementia Rating Scale

This study was conducted in accordance with the declaration of Helsinki and was approved by the Rennes University Hospital ethics committee (approval number IDRCB: 2011-A00392-39). After a complete description of the study, all patients gave their informed written consent.

### 2.2. Experimental task

Patients were asked to perform a motivated Simon task developed in our team (Houvenaghel et al., 2016a, b). Each trial began with the display of a monetary incentive cue in the form of a coin that was either 1 cent, 1 € or fake (a chimeric combination of a 1 cent and 1€ coin, named fake stimulus in the following sections). Each coin had a diameter of 3.7 cm that subtended a visual angle of 2.35°. After a black screen (700-1100 ms), a blue or yellow circle appeared either on the right or on the left side of the screen. Each circle had a diameter of 5.5 cm that subtended a visual angle of 2.48°. Participants had to press a blue or a yellow button on a keyboard with the left or right hand as fast and as accurately as possible according to the color of the stimulus while ignoring its location. The left and right color positions were counterbalanced across subjects. Two different conditions were possible: a congruent one, when color and location matched and activated the same response; and an incongruent one, when color and location indicated different responses, which induced response conflict. In order to obtain the full reward, the participants had to respond as fast and as accurately as possible, with the reward size being proportional to speed. Baseline reaction time (RT) was calculated in the practice phase (32 trials without a reward cue), and participants actually had to be one sixth faster than their baseline mean RT to obtain the full reward. A performance equal to baseline yielded only 50% of the promised reward, while slower and erroneous responses were not rewarded. At the end of each trial, a feedback showing the amount of money virtually won since the beginning of the block was displayed for 1500 ms before the next trial. The total virtual money won was also displayed at the end of each block of trials. Both baseline and experimental phases (5 blocks of 72 trials) contained the same amount of congruent and incongruent trials, and each congruence/level of reward combination was displayed 60 times in a pseudo-randomized order.

### 2.3. Recordings

All patients were studied under antiparkinsonian medication, two days after surgery and before subsequent implantation of the subcutaneous stimulator. STN LFPs were recorded through a g.BSamp^®^ (g.tec Medical Engineering, Schiedlberg, Austria) biosignal amplifier connected to a PowerLab^®^ 16/35 (ADInstruments, Dunedin, New Zealand) system. Intracranial activity was recorded bipolarly from all two adjacent contact pairs of each DBS electrode. Thus, for each electrode, three bipolar derivations were recorded: 0-1, 1-2, and 2-3. Signals were amplified and sampled at 1000 Hz and monitored online using the Labchart^®^ software (ADInstruments). Triggers corresponding to the task stimuli were recorded simultaneously.

### 2.4 Behavioral analyses

Behavioral data analyses were performed using R (version 3.4.2; R core team, 2017) implemented with the nlme (Pinheiro et al., 2014) and lme4 (Bates et al., 2015) packages. Only the experimental phase trials with RT comprised between 200 ms and 1500 ms were analyzed. Trials slower than 3 standard deviations from the mean were also excluded. This resulted in the removal of 1.5 % of the dataset. RT and accuracy were analyzed as a function of congruence and motivation.

Statistical analyses consisted in comparing RT and accuracy according to congruence and the level of motivation. This resulted in 2 (congruence) * 3 (reward cue) factorial designs, with two levels of congruence (congruent and incongruent) and three levels of reward cue (fake, 1 cent, 1€). To perform these analyses, we used linear (using the lme function) and non-linear mixed models (using the glmer function) on RT and accuracy, respectively. Since RT were not normally distributed, they were inverse-transformed before the analysis. We used mixed-models instead of standard ANOVAs to avoid the loss of power associated with averaging data and to take into account inter-individual variability by adding a random effect of subject (see Gueorguieva and Krystal, 2004). Post-hoc analyses were carried-out when main effects were significant. We used Tukey tests computed by the glht function (package multcomp) providing adjusted p-values using individual *z* tests (Hothorn et al., 2007). In the case of significant congruence*motivation interaction on RT or accuracy, further models were ran between each pair of motivation condition to evaluate the differences in the congruence effect between the two motivation conditions. In those cases, p-values were adjusted using the Bonferroni correction. The significance statistical threshold was set at p = 0.05.

### 2.5 LFPs signal analyses

Signal preprocessing was performed using the Brainstorm toolbox (Tadel et al., 2011) for Matlab (The Mathworks, USA). All subsequent data analyses were performed using custom Matlab code (available here [insert URL]) based on published equations (Cohen, 2014).

#### 2.5.1 Preprocessing

All LFP data was high-pass filtered offline at 0.5 Hz. Data was epoched from -1 to 2s surrounding the presentation of the reward cue, and also surrounding the imperative stimulus. Thus, two sets of epochs were analyzed: one to investigate LFP responses to the motivational cue, and one to investigate LFP responses to the imperative stimulus. Such long epochs were used on purpose to avoid edge artifacts associated with wavelet convolution (see next section). All trials were visually inspected, and those with excessive noise or artifacts were manually discarded. Subsequent analyses only focused on correct trials with respect to the behavioral preprocessing steps described above. After these preprocessing steps, there were an average of 99 (± 20) and 47 (± 10) trials per condition for each STN in the motivation (3 reward conditions) and cognitive action control (3 [reward] * 2 [congruence] conditions) epoching respectively. As a result of excessive noise in most trials, one STN data was completely discarded in two patients, thus resulting in 30 analyzable STN datasets.

The most distal contacts were located 11.5 ± 4.2 mm lateral to the anterior-posterior commissure (AC-PC) line, -3.1 ± 1.7 mm posterior to the middle of the AC-PC line, and -5.4 ± 1.7 mm below this point. These positions and the CT-scan confirmation of the anatomical location of the lead indicated that the pair of distal contacts (0-1) was effectively located within the STN. We thus focused the subsequent analyses on the most distal pair of contacts.

#### 2.5.2 Time-frequency analyses

Time-frequency decomposition of data was performed using a complex Morlet wavelet convolution in the frequency domain. The power spectrum of the fast-Fourier transform of the LFP signal was multiplied by the power spectrum of a set of complex Morlet wavelet 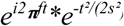, where t is time, f is frequency ranging from 1 to 40 Hz in 50 logarithmically spaced steps, and s is the width of each frequency band defined by *n*/(2π*f*) (using a logarithmically increasing n from 4 to 10). The inverse fast-Fourier transform was then applied to the result to obtain the analytic signal. Power was then extracted from the resulting complex signal by taking the squared magnitude of the signal at each time point (real[z(t)]^2^ + imaginary[z(t)]^2^). Power was then normalized using a decibel (dB) transform [dB power = 10 × log 10 (power/baseline)] to ensure that all data were at the same scale, thus allowing condition comparisons. Since we wanted to investigate the effect of congruence and motivation and their interaction on the LFPs, we used a baseline of -500 to -200 ms before the onset of the reward cue for both the motivation and cognitive action control sets of epochs. Average baseline power was calculated across all conditions using that time-window.

We also investigated phase results by computing inter-trial phase clustering (ITPC) to investigate local functional organization. This measure shows similarity regarding the phase angle of oscillations across trials, thus indicating the level of functional organization. It was computed for each time-frequency point according to 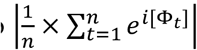 where n is the number of trials, and Φ is the phase angle at trial t. This formula calculates the length of the average phase angle complex vector, and takes values ranging from 0 (indicating a uniform phase angles distribution across trials) to 1 (indicating perfect clustering of phase angles). To take into account differences in trial count between conditions, and to ensure that ITPC would be comparable, we transformed ITPC to ITPCz as follows: ITPCz = n x ITPC^2^ where n is the number of trials (see Cohen, 2014). ITPCz is expressed in arbitrary units (a.u).

Left and right STN were assumed to be independent (as in Zavala et al., 2013; 2014), thus analyses were applied to 30 pairs of contacts (2 STN data were entirely discarded, see preprocessing section). Power and ITPC were compared across conditions, and for both motivation and cognitive action control epochings, by visually selecting time-frequency windows of interest on the condition and subject average time-frequency map (downsampled to 100 Hz). We chose this approach rather than the classic permutation testing because of the factorial nature of our experimental design (see Cohen, 2014). Note that selecting windows of interest on average maps is orthogonal to the effect of interest, and thus not subjected to circular inference. To better consider inter-individual variability, smaller subject-specific time-frequency windows were defined within the first window of interest. These subject-specific windows were centered around the time-frequency point of maximum power/ITPCz for each subject. The size of those smaller windows was of 3 frequencies (according to the logarithmic scale) by 13 time points (so around 77 ms after downsampling). Power and ITPCz values were then extracted from those windows and subjected to further statistical analyses.

For each time-frequency window, and for both the motivation and cognitive action control sets of epochs, power and ITPCz were compared between conditions using the same linear mixed models as in the behavioral analysis. This resulted in a 3 (reward cue) and 3 (reward cue) * 2 (congruence) factorial design for the motivation and cognitive action control sets of epochs respectively. For each analysis, the Bonferroni corrections was applied to account for the number of time-frequency windows selected so that the threshold was p = 0.05/n (time-frequency windows). When main effects were significant, post-hoc Tukey test were applied as described above.

## 3 Results

### 3.1 Behavioral results

The typical congruence effect was reflected by a strong slowing effect of the incongruent stimulus observed on RT (F(1, 6189) = 302.1, p<0.0001; Figure 2A, Table 2). The promised reward also significantly affected RT (F(2, 6189) = 103.2, p<0.0001), since we observed faster responses for the 1€ condition compared to the 1c and fake reward conditions (1€ vs. 1c, p<0.0001; 1€ vs. fake, p<0.0001; 1c vs. fake, p = 0.08). These results illustrate that patients were faster to respond when the high reward cue was presented. Conflict resolution was also influenced by the promised reward: the effect of congruence was different according to the motivation condition as revealed by a significant congruence*motivation interaction (F(2, 6189) = 6.03, p = 0.002). In order to test which reward condition had the strongest congruence effect, we ran again the same model by isolating each pair of reward conditions. Since three different models were ran, we applied a corrected significance threshold of p = 0.05/3 = 0.017. Only the comparison between 1€ and fake yielded a significant congruence*reward interaction, showing that the congruence effect was higher, and conflict resolution harder, for the 1€ cue as compared to the fake cue (60 vs. 35 ms; F(1, 4127) = 11.9, p<0.0001; all other comparisons NS). This higher congruence effect in 1€ condition seemed mostly driven by the faster congruent RT. Indeed, the congruent condition for the 1 € reward had an averaged RT of 470 ms compared to 515 and 531 ms for the 1c and fake rewards, respectively; thus a difference of 45 and 61 ms, respectively (all multiple comparisons between rewards for the congruent conditions p<0.0001). For the incongruent conditions, the multiple comparisons were also all significant (p<0.0001), except between the 1c vs. fake conditions (p=0.05). RT differences were significantly less pronounced in this condition, with faster RT still observed for the 1€ condition. Average RT for 1c and fake were of 552 ms and 566 ms conditions, leading to a difference of 16 and 39 ms with the 1€ condition (average RT: 527 ms) respectively.

**Figure 1:**
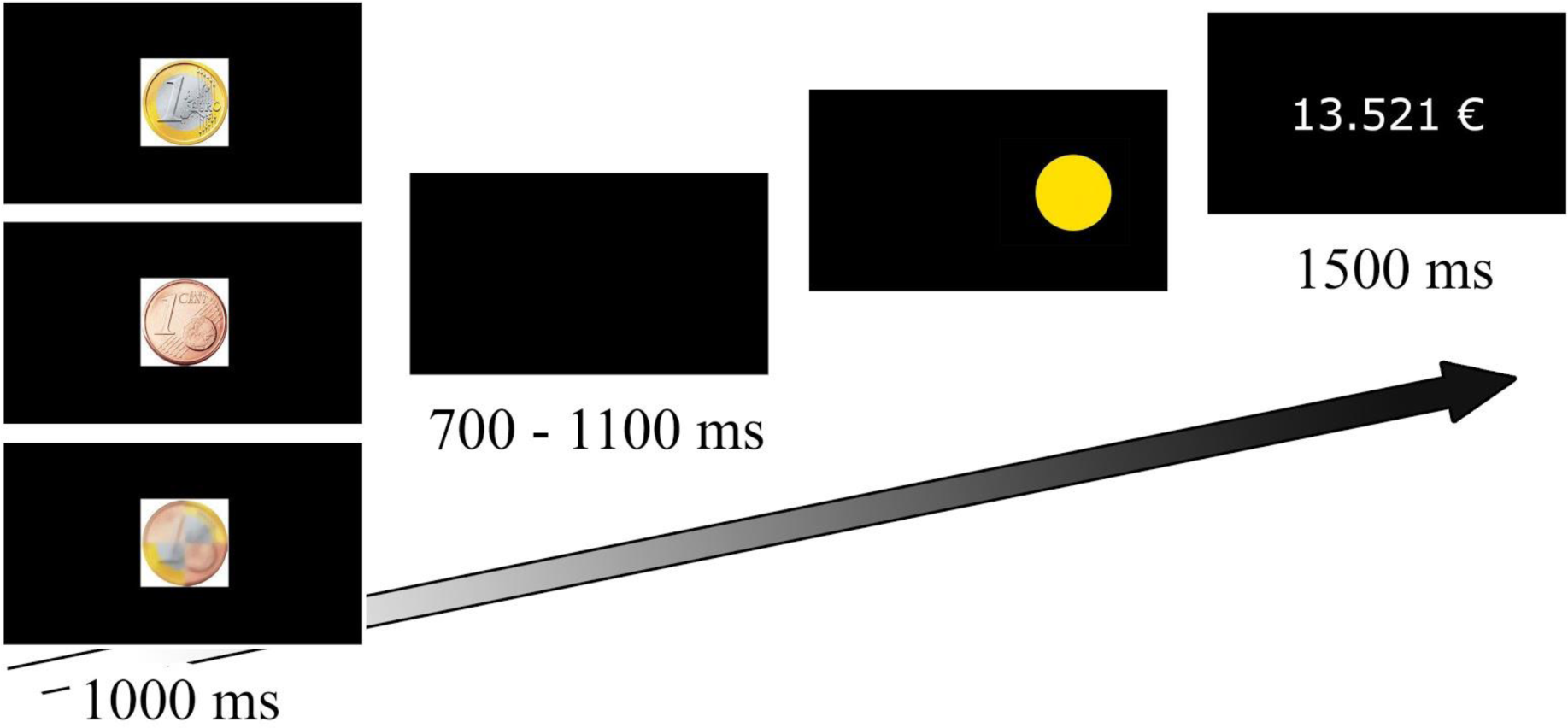
At the beginning of each trial, the incentive monetary cue (€0, 1 cent or €1) was displayed for 1000 ms. After an interval of 700-1100 ms, a blue or yellow circle appeared on the left- or right hand side of the screen, and remained there until participants pressed the blue or yellow button of the keyboard (as quickly and accurately as possible). At the end of each trial, the size of the reward accumulated since the start of the relevant experimental block was displayed for 1500 ms.

**Figure 2:**
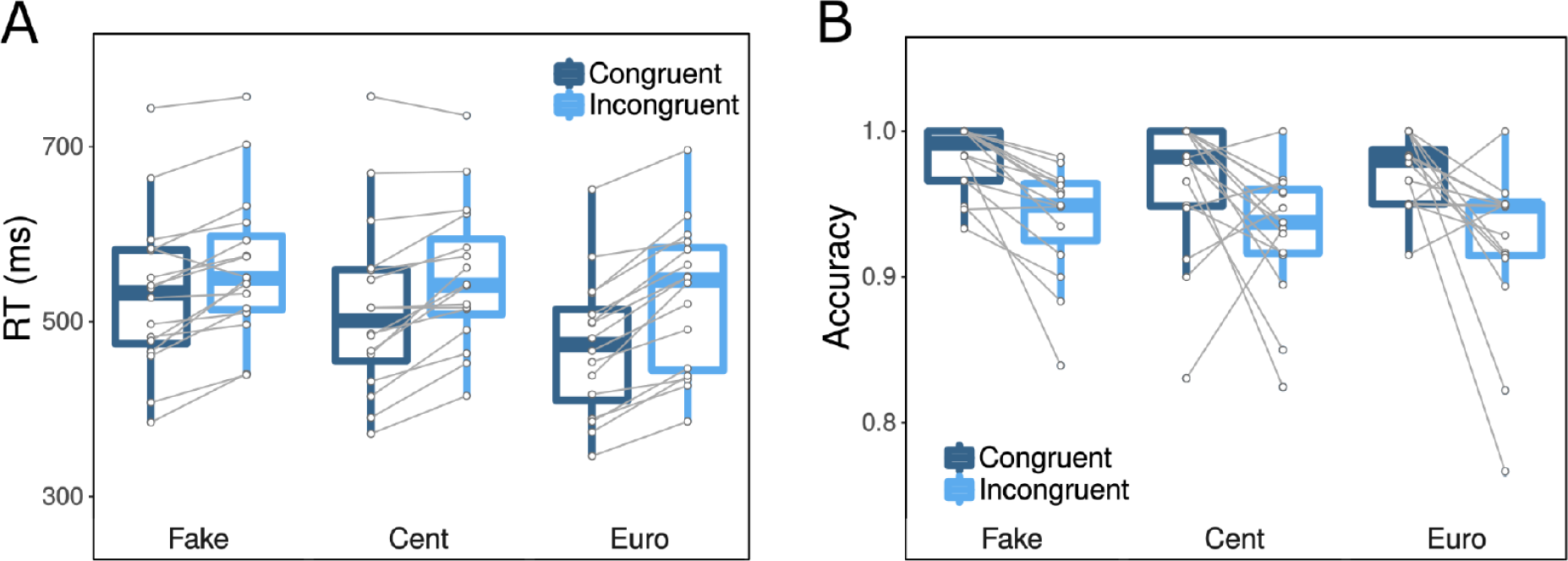
Box plots of RT (A) and accuracy (B) as a function of congruence and size of the promised reward. Overlaid data points show average RT (A) and accuracy (B) for each patient.

**Table 2:**
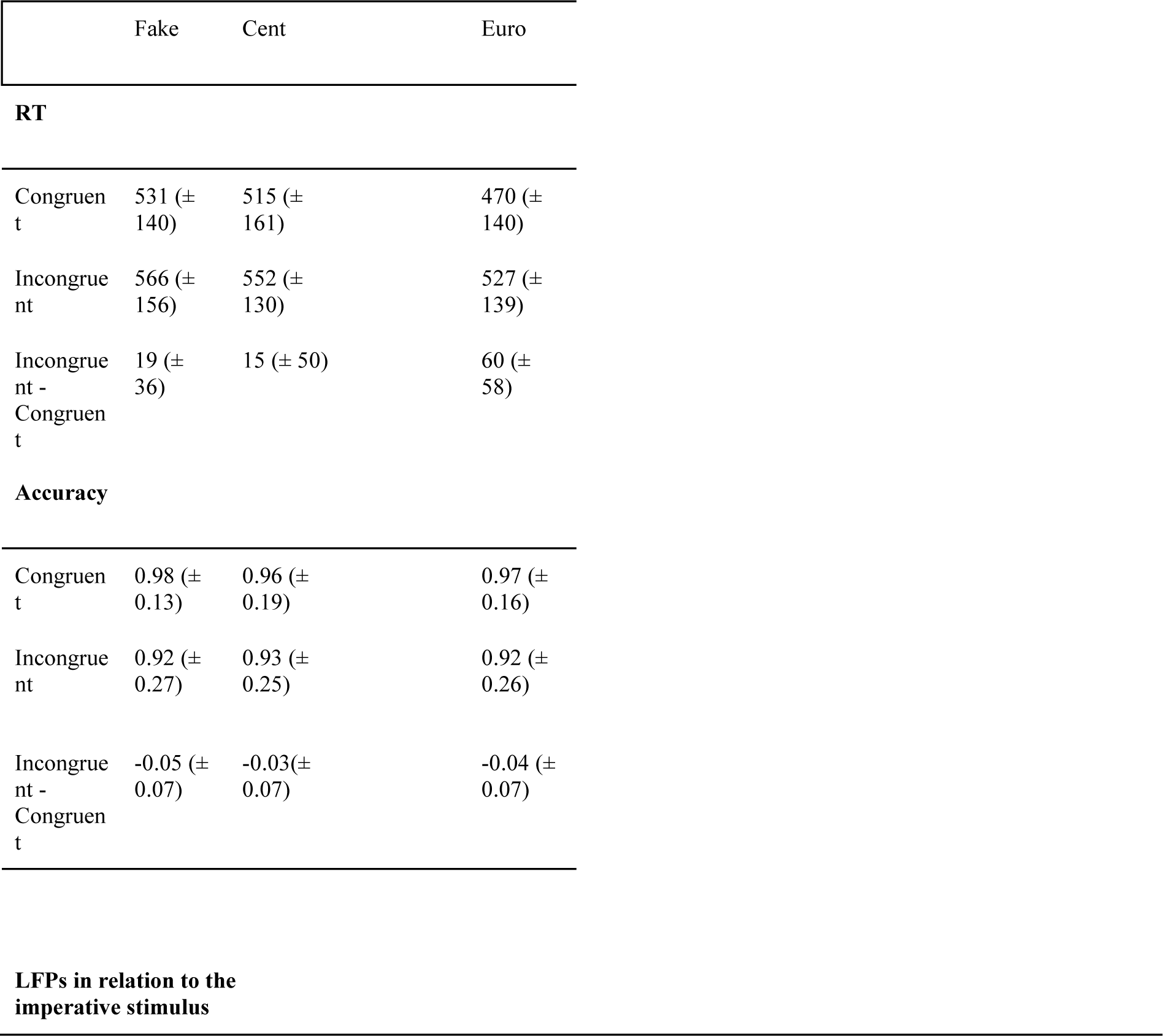
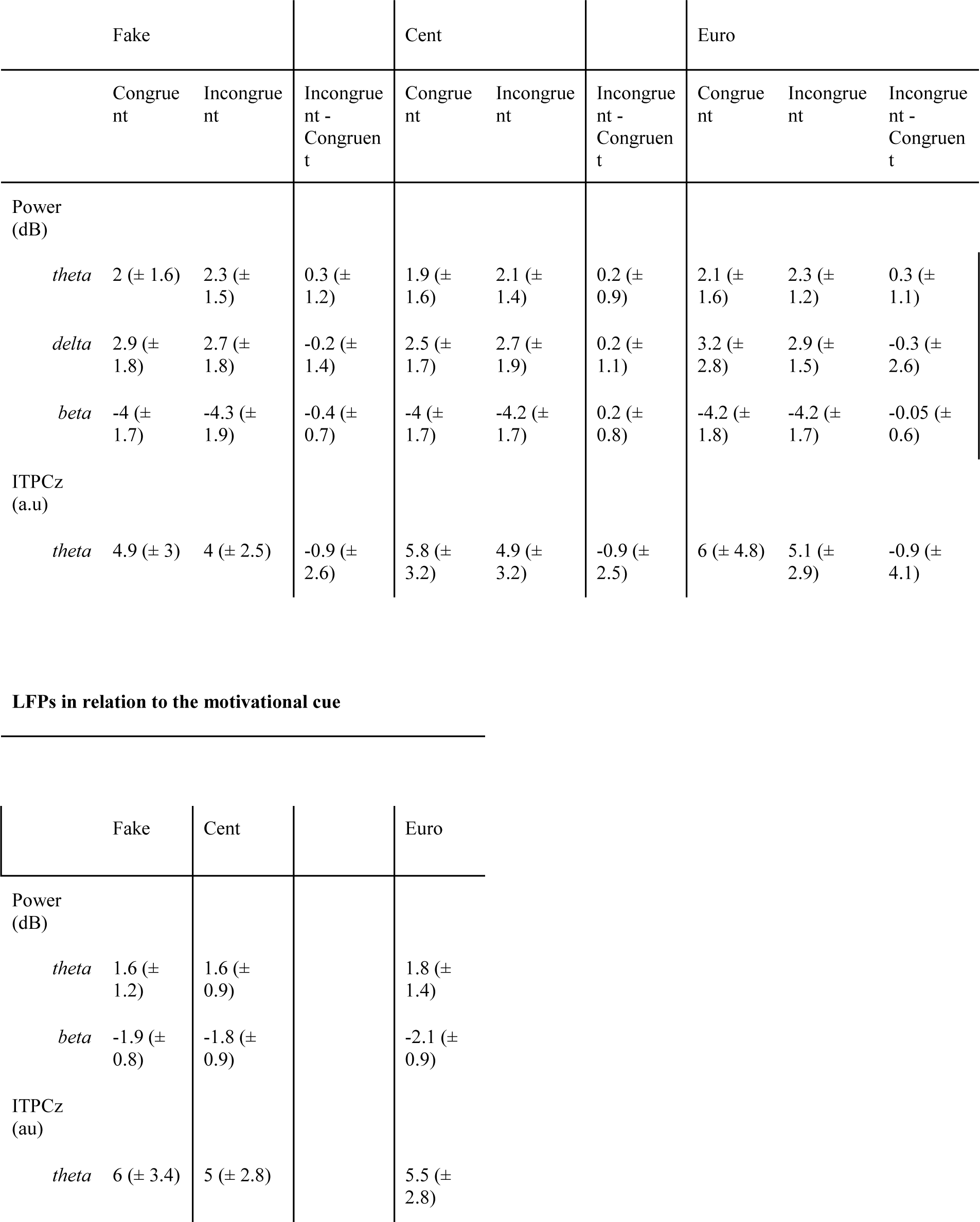
Behavioral and LFPs results showing mean (SD) RT and accuracy, as well as power and ITPCz extracted from relevant time-frequency windows, according to congruence and reward size.

As for RT, a strong congruence effect was also observed for accuracy, with less accurate responses in the incongruent condition (χ^2^ = 67.5, p<0.0001, Figure 2B, Table 2). However, contrary to RT, accuracy was not different according to the size of the promised reward (χ^2^ = 0.4, p=0.8). The congruence effect was also not influenced by motivation, as revealed by the absence of a significant congruence*motivation interaction (χ^2^ = 2.8, p=0.2).

### 3.2 LFPs in relation to the motivational cue

In this section, we focused on testing if the STN had specific responses according to motivation. Therefore, we investigated whether LFPs fluctuated according to the size of the promised reward prior to the presentation of the imperative stimulus.

#### 3.2.1 Power results

For this set of analyses, we focused on 2 different time-frequency windows (Figure 3A): a first one in the theta-lower alpha band (from 5.2 to 10.3 Hz and from 100 to 450 ms after the reward cue), and a second one in the beta band (from 12 to 23.5 Hz and from 200 to 650 ms). Any activity occurring after 1700 ms was not considered for analyses, since the imperative stimulus could be displayed between 1700 and 2000 ms. Furthermore, we did not consider the delta increase around 700 ms, since it was already present before stimulus onset, as well as the late theta suppression because it is hardly dissociable from an effect that would occur around the imperative stimulus onset. Since 2 time-frequency windows were selected, the significance threshold was adjusted to p = 0.05/2 = 0.025.

**Figure 3:**
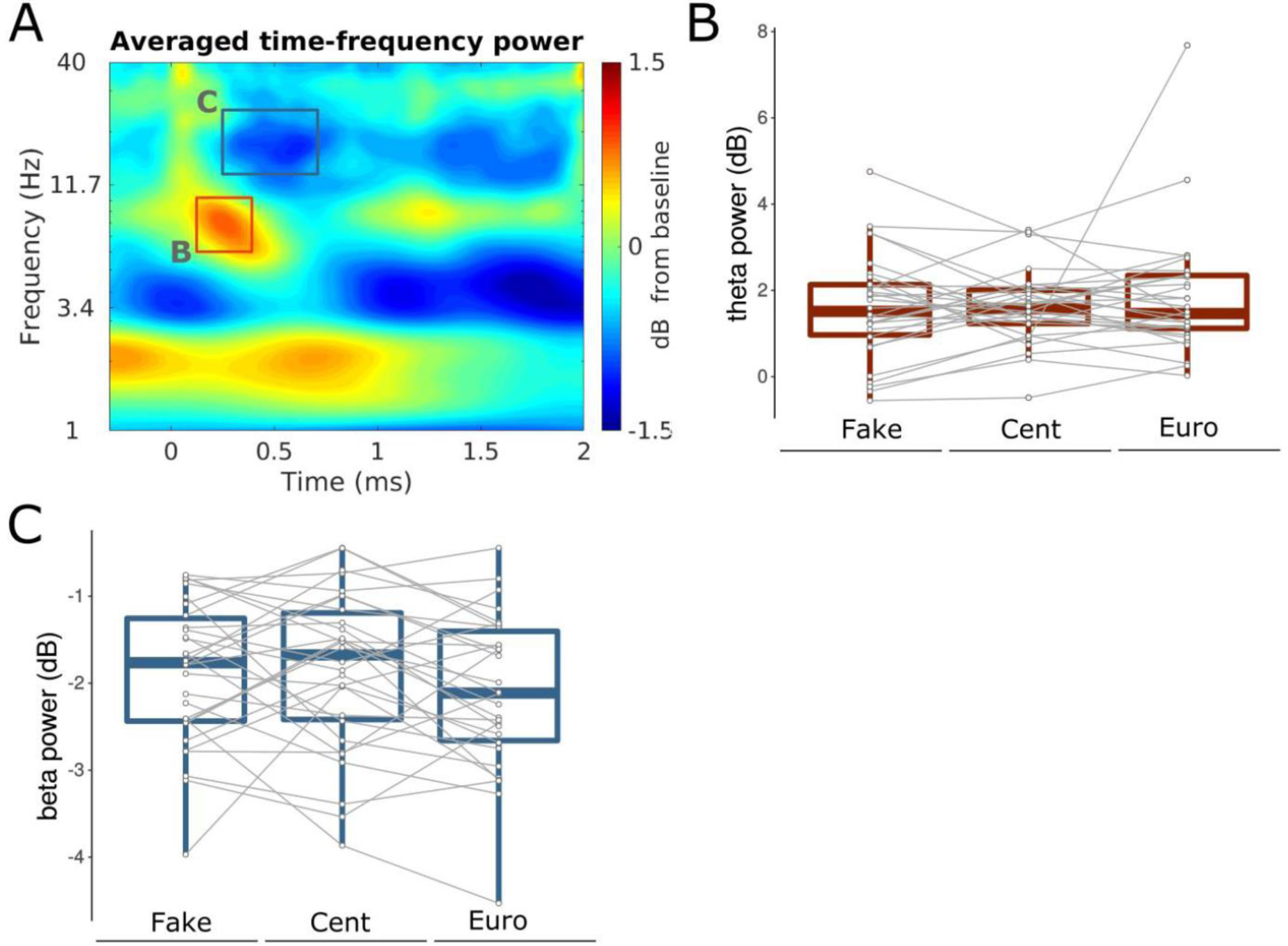
Panel A: Average time-frequency power in relation to the reward cue. Time 0 corresponds to the onset of the cue. The squares correspond to the time-frequency windows chosen for the analyses in the theta (B) and beta (C) band. Panels B and C: Boxplots of the changes in theta (B), and beta (C) power (dB) according to the size of the promised reward. Overlaid data points show the average power for each STN.

##### a. Theta-lower alpha window

We observed an increase in theta power as compared to baseline, that peaked at an average of 7.6 ± 1.6 Hz and 296 ± 116 ms. Figure 3B illustrates the power extracted around that peak for each reward cue. We observed no clear fluctuation in theta power according to the size of the promised reward, which was confirmed by the absence of a significant motivation effect (F(2, 58) = 0.5, p = 0.6; Figure 3A, Table 2). This result suggests that the amplitude of the STN oscillatory activity was similar regardless of motivation.

##### b. Beta window

We observed a beta power decrease as compared to baseline, that peaked at an average of 16.8 ± 3.9 Hz and 497 ± 142 ms. Figure 3C presents the power extracted around that peak for each motivational cue and shows that the decrease in power was stronger for the highest motivation level (F(2, 58) = 4.2, p = 0.02; Table 2). Post-hoc tests confirmed that the decrease was stronger for the 1€ condition as compared to the 1c condition (p = 0.04) and to the fake condition (p = 0.03). There was no difference between the 1c and fake conditions (p = 1).

#### 3.2.2 ITPC results

Figure 4A presents the averaged ITPCz time-frequency map where higher phase alignment can be observed in the theta band. We selected a time-frequency window ranging from 2.3 to 7 Hz and from 130 to 690 ms. This window roughly corresponds to the theta window investigated for power analyses. Later ITPCz fluctuations were not considered since the imperative stimulus could be presented beginning at 1700 ms after presentation of the reward cue which prevents interpretations.

**Figure 4:**
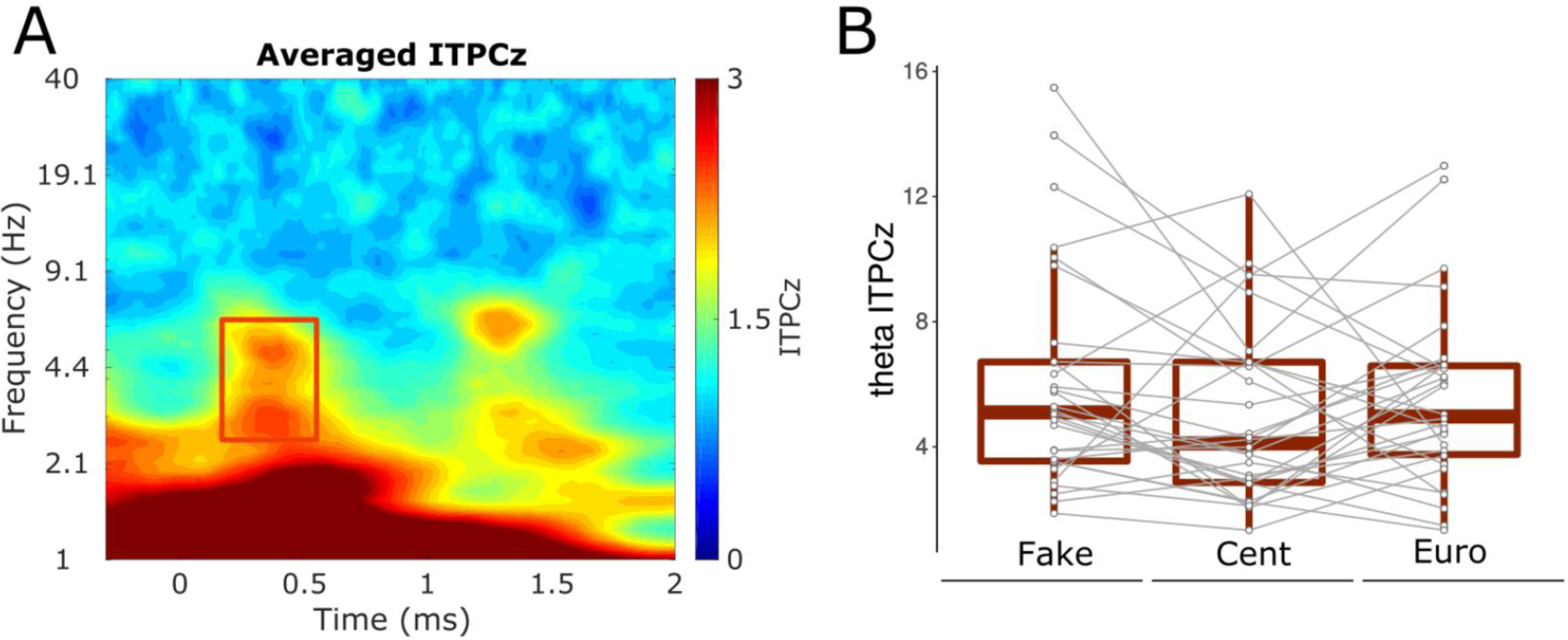
Panel A: Average time-frequency ITPCz in relation to the reward cue. Time 0 corresponds to the onset of the cue. The square corresponds to the time-frequency windows chosen for the analyses in the theta (B) band. Panel B: Boxplots of the changes in theta ITPCz according to the size of the promised reward. Overlaid data points show average ITPCz for each STN.

In the selected window, ITPCz peaked at 4.3 ± 1.5 Hz and 387 ± 171 ms. Although phase alignment seemed to be higher in the fake condition, no effect of motivation was found on ITPCz (F(2,58) = 1.5, p = 0.2; Figure 4B, Table 2). This suggest that the functional organization of theta oscillations was similar across the different possible sizes of the promised reward.

### 3.3 LFPs response to the imperative stimulus

Here, we wanted to test if STN oscillatory activity was modulated by the need for cognitive action control, and if this modulation was motivation-specific. We thus investigated if time-frequency power and phase clustering fluctuated according to conflict and the size of the promised reward.

#### 3.3.1 Power results

Figure 5A presents the cross-condition average time-frequency map of power. We selected 3 different time-frequency windows for the investigation of the effect of conflict and motivation on power: a theta window (from 100 to 400 ms and from 5.2 to 10.3 Hz), a delta window (from 350 to 1310 ms and from 1 to 3.3 Hz) and an upper alpha-beta window (from 100 to 1100 ms and from 12 to 34.4 Hz). Inside those windows, smaller subject-specific windows around power peak were defined. Since 3 time-frequency windows were selected, the significance threshold was adjusted accordingly using the Bonferroni correction to p = 0.05/3 = 0.017.

**Figure 5:**
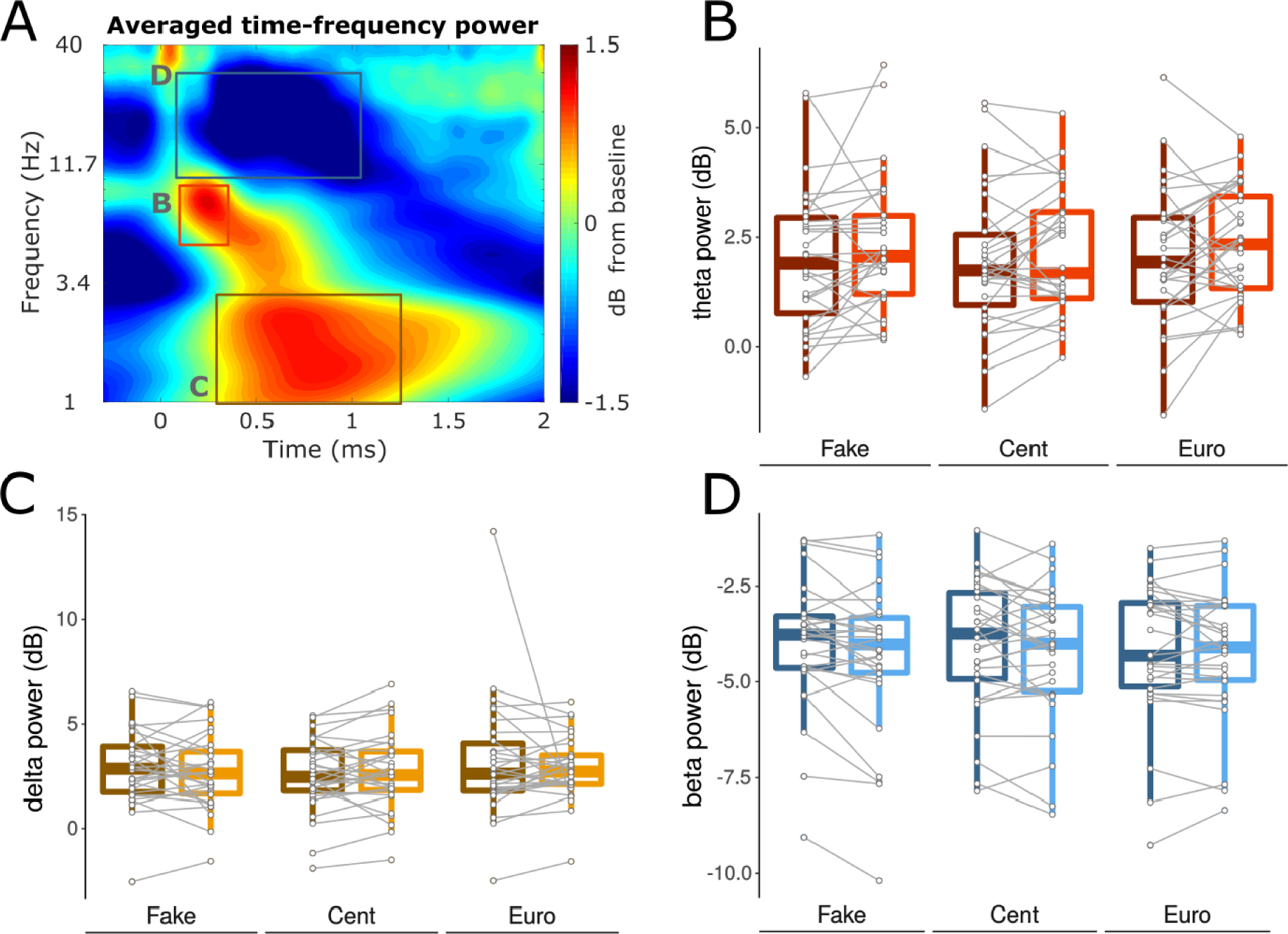
Panel A: Average time-frequency power plot (relative to baseline, in dB) for the imperative stimulus. Time 0 corresponds to the stimulus onset. The squares correspond to the time-frequency windows chosen for the analyses in the theta (Panel B), delta (Panel C), and beta (Panel D) band. Panels B-D: Boxplots of the changes in theta (B), delta (C), and beta (D) power (dB) according to congruence and size of the promised reward. Overlaid data points show average power for each STN.

##### a. Theta window

We observed an early increase in power from baseline in the theta window, with an average peak at 7.4 ± 1.6 Hz and 290 ± 100 ms. The extracted theta power was influenced by conflict (Figure 5B, Table 2), since we observed a higher power increase in the incongruent condition as compared to the congruent one, as revealed by a significant congruence effect (F(1, 145) = 6.0, p = 0.01). However, the size of the promised reward had no effect on theta power (F(2, 145) = 0.9, p =0.4). In the same way, reward conditions had no effect on the size of the congruence effect, as revealed by the absence of a significant motivation*congruence interaction effect (F (2, 145) = 0.1, p = 0.9).

##### b. Delta window

Increased power from baseline could also be seen later in time in the delta band, with an average peak of power at 2.3 ± 1 Hz and 835 ± 284 ms. We did not observe any differential increase in power according to congruence (F(1, 145) = 0.3, p = 0.5; Figure 5C, Table 2), or according to the size of the promised reward (F(2, 145) = 1.5, p = 0.2). The absence of a significant congruence*motivation interaction further showed that the congruence effect was similar across the motivation conditions (F(2, 145) = 0.7, p = 0.5).

##### c. Upper alpha-beta window

Power extracted from the upper-alpha-beta window actually peaked in beta with an average peak at 18.3 ± 5.2 Hz and 596 ± 267 ms. The beta power decrease from baseline seemed to depend on congruence with stronger decrease in the incongruent condition. However, this effect was not significant using the statistical threshold corrected for multiple comparisons (F(1, 145) = 5.2, p = 0.02; Figure 5D, Table 2). Beta band power was not affected by the size of the promised reward (F(2, 145) = 0.4, p = 0.7),nor did we observe a significant influence of motivation on the size of the congruence effect (F(2, 145) = 1.1, p = 0.3).

#### 3.3.2 ITPC results

We further investigated the functional organization of neural oscillations across trials. To this end, we computed ITPCz at all time-frequency points. Similarly to power, we focused the analyses of the experimental condition effects on a specific time-frequency window. Here, we selected only one window with an increase in ITPCz in the theta band from 2.3 to 8.2 Hz, and from 80 to 460 ms (Figure 4A). This increased activity peaked at 5.2 ± 1.8 Hz and 274 ± 127 ms.

As shown in Figure 6B, ITPCz was consistently lower when the situation was incongruent (F(1, 145) = 7, p = 0.01; Table 2), indicating that the conflict induced a lower functional organization compared to congruent situations. As opposed to power analyses, ITPCz varied according to motivation with functional organization in theta increasing with the size of the promised reward. This was revealed by a significant motivation effect (F(2, 145) = 4.3, p = 0.01). However, the size of the congruence effect was not affected by motivation and was similar across the possible promised reward sizes (F(2, 145) = 0, p =0.9).

**Figure 6:**
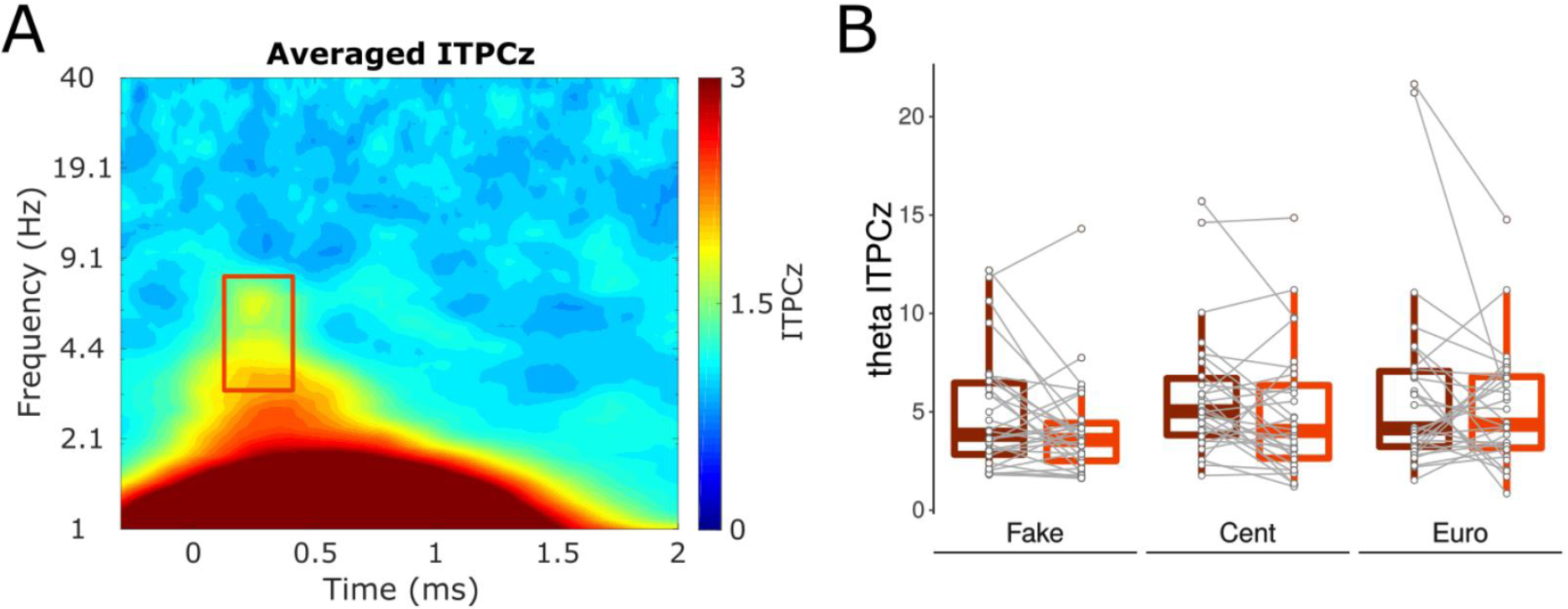
Panel A: Average time-frequency ITPCz in relation to the imperative stimulus. Time 0 corresponds to the onset of the stimulus. The square corresponds to the time-frequency window chosen for the analyses in the theta band. Panel B: Boxplots of the changes in theta ITPCz according to congruence and size of the promised reward. Overlaid data points show average ITPCz for each STN.

### 3.4 LFPs-behavior relationships

Here, we tested whether STN oscillatory activity, in terms of power, was linked with behavior. First, since beta power following the reward cue showed a significant effect of reward, and since reward had a significant effect on RT, we checked whether these effects correlated with each other. No significant correlation was observed between the reward effect on RT and beta power, neither on the 1€-fake reward effect (ρ = 0.26, p = 0.16), nor the 1€-1cent effect (ρ = 0.11, p = 0.55). We did not identify any other significant correlation between power in other time-frequency windows and behavior.

## Discussion

The goal of this study was to improve our understanding of the role of the STN during incentive motivated action control. To this aim, we analyzed STN oscillatory activity in response to promised-reward during conflict resolution. Indeed, although the influence of reward expectation on cognitive action control has yielded somewhat different results so far, most of them have suggested an influence of the reward presentation on conflict resolution. Since conflict modulates low-frequency power in the STN and since these oscillatory dynamics seem to fluctuate according to reward stimuli, we hypothesized that promised-reward during a conflict task would modulate the size of the congruence effect on STN for low-frequency oscillations. Although our result did not confirm this hypothesis, we identified subtle reward effect on STN activity during conflict resolution.

## Promised-reward toughens conflict resolution

Our results illustrate that the size of a promised-reward influences cognitive action control by increasing the size of the congruence effect on RT. In other words, high rewards were associated with increased difficulties in solving conflict. Although these results differ from studies, showing a beneficial (Padmala & Pessoa, 2011) or no effect (Aarts et al., 2014; van den Berg et al., 2014) of reward presentation, they are in line with other studies associating cognitive action control difficulties with reward presentation (Padmala & Pessoa, 2010; Houvenaghel et al., 2016a). In Houvenaghel et al. (2016a; 2016b), the same experimental paradigm was used and revealed that cognitive action control was also less efficient with higher reward. However, in PD patients this effect appeared only when considering the dynamics of impulsive action suppression in Houvenaghel et al. (2016a), while it was already present at the average RT level in Houvenaghel et al. (2016b). Although these results suggest an impact of promised reward on conflict resolution, the fact that other studies found different, or no effects, and that results vary using the same experimental task, point to a subtle reward influence that might be hard to interpret in the light of the sole behavioral results.

## Promised-reward modulates the functional organization of low-frequency STN activity during conflict resolution

The STN holds a crucial position in the cortical-subcortical loops, since it receives many inputs from circuits that are involved in cognitive action control and in reward processing (Isaacs et al., 2018). As a result, the STN activity might reflect integration of both cognitive processes. This proposition has received support from recordings of STN activity in animals (Isoda & Hikosaka, 2008; Espinosa-Parilla et al., 2015) and in humans (Zaghloul et al., 2012; Rosa et al., 2013; Fumagalli et al., 2015).

Conflict has been associated with increase in theta band power in STN LFPs in numerous studies (see Zavala et al., 2015 for a review). In line, we also report here a robust increase in theta power, as well as a stronger beta power decrease, when the situation was incongruent as compared to congruent. Although increased theta activity has been consistently associated with conflict resolution, beta activity has been proposed to be a signal of response inhibition. So far, whether beta suppression or enhancement is involved in inhibition is still a matter of debate (see Zavala et al., 2015). However, one could interpret our results in line with Yugeta et al. (2013), claiming that stronger beta suppression in antisaccade trials would be linked to a stronger inhibition of the automatic response yielded by the side of the stimulus presentation.

Local functional organization, inferred through ITPC, was also influenced by conflict, but in an opposite direction to what was expected, with higher theta ITPC during congruent as compared to incongruent trials. Although this result has already been reported in a previous study (Zavala et al., 2013), since conflict typically yields increased theta power, higher functional organization would be logically expected. One explanation of such a result might be found in the usually higher variability observed in incongruent RT. Indeed, congruent RT had less variance than incongruent RT, which in turn implies that all response-locked oscillatory activity had a greater chance to cluster across trials in congruent versus incongruent trials. This could be verified by applying response-locked analyses to the present data set. However, due to technical difficulties, RT could not be associated to triggers in the present LFP signals and were not associated with a trial number in the recordings, preventing such analyses from being conducted.

As opposed to our hypothesis, we did not observe any modulation of conflict-related theta power increase according to the size of the promised reward, nor did we find any global reward effect on STN LFP power following the imperative cue. However, a reward effect was present when focusing on theta ITPC, illustrating that high rewards was associated with greater local functional organization during conflict resolution. Although reward did not influence the congruence effect on theta ITPC, to our knowledge, this is the first empirical evidence that reward stimuli modulates STN activity during conflict resolution. This result suggests that STN low-frequency activity, especially in the theta band, is involved in reward processing during cognitive action control. Our results reveal subtle reward effects, but are in line with the fact that the STN is a convergent site for cortical areas involved in both processes (Maurice et al., 1998; Isaacs et al., 2018). The subtle nature of these effects are also in line with the behavioral ones that reveal a varying influence of reward according to the different studies (Padmala & Pessoa, 2011; Aarts et al., 2014; van den Berg et al., 2014; Padmala & Pessoa, 2010; Houvenaghel et al., 2016a, 2016b). Elucidating the exact mechanisms through which STN low-frequencies participate in using reward information during the information processing required for conflict-resolution is beyond the scope of this study, but the results presented here encourage for future investigations in that direction.

## Reward presentation influences high rather than low-frequency STN activity

Our results did not reveal any effect of the reward size on theta power after the onset of the reward cue. This is at odd with the results from Zénon et al. (2016), who described increased low frequency power with higher reward. In their study, decision conflict was inferred on the basis of the probability to accept/refuse performing a trial given the size of the reward and the amount of effort to provide. This is different, as the authors explained, from classical conflict tasks with a two alternative forced choice. However, although it is hard to compare both studies in terms of conflict-related activity, modulation of STN activity in relation to the reward cue should be similar. The absence of reward presentation effect could be related to a potential anticipation of conflict. Indeed, Cooper et al. (2017) have described a strong fronto-parietal increase in theta power in anticipation of conflict that could predict cognitive action control performances. Although this study was performed in healthy participants, since the STN has been associated with cognitive action control performance, such anticipatory theta activity might also occur in the STN in the framework of proactive cognitive action control. This could in turn mask subtle reward effects and one could argue that, since response conflict was much less prominent in Zénon et al. (2016), these anticipatory oscillatory changes were less present, thus allowing for the detection of reward-related fluctuations in power. Although this is purely speculative, it could also explain why no reward effect was found in low-frequency activity in Oswal et al. (2013), in which response conflict (when a limb was cued and a movement of an opposite limb was expected) was high.

As opposed to low frequencies, STN activity in higher frequencies following cue onset, specifically in the beta band, was influenced by the size of the reward. More precisely, higher rewards were associated with stronger beta power suppression. As it was argued in the previous paragraph, interpretation of beta power suppression is challenging. This suppression, taking place before the imperative stimulus was displayed, might reflect the preparation for the upcoming action, and could also indicate response withholding during the pre-stimulus period. Interestingly, higher reward also had an effect on behavior by speeding RT, which would be in line with a response preparation sustained by beta suppression. However, we did not find any relationship between these behavioral results and STN beta activity, thus encouraging to interpret this result with caution.

As a whole, although our results show a reward effect in the STN oscillatory activity, it is different from what has already been observed. One explanation could be that experimental designs, in terms of experimental task, differed. An interesting fact related to the heterogeneity in STN activity modulation facing motivation is that researches also pointed out that there is also heterogeneity in how STN neurons respond to reward (see Bonnevie & Zaghloul, 2018). Although a direct link with the present results is far-stretched, further investigations could focus on the contributions of these differential neuronal responses to the STN LFPs signal.

## Limitations

This study comes with two major limitations. First, we did not observe any significant relationship between behavior and STN neuronal activity. Thus, although we found changes in STN activity related to task conditions, the absence of correlation with behavioral results makes difficult to interpret electrophysiological findings as relevant for behavior. On the other hand, we believe that this limitation might result from the second limitation of this study, which is linked to technical difficulties in having RT corresponding triggers in the LFP signal. Indeed, correlations between STN activity and behavior were estimated using mean RT and power results while the benefit of investigating single trial brain-behavior relationships has been demonstrated (Cohen & Cavanagh, 2011). Thus, although a true relationship might exist between these measures, we were unable to expose it since we couldn’t use each trial RT-power datapoint. Especially, this would explain why we did not find any relationship between theta power and RT, which has already been described in the STN (Zavala et al., 2013), and repeatedly observed in the midfrontal cortex (see Cohen, 2014 for a review). Further studies will have to deal with the issue of relationships between the effect of reward on behavior and STN activity.

## Conclusion

The STN is a key structure in the basal ganglia, since it shares hyperdirect connections with cortical areas that have been described as essential in various cognitive processes, notably in cognitive action control and reward processing. Recent research have proposed that incentive motivation influences cognitive action control performances. This study provides empirical evidence that STN activity, as behavior, is influenced by reward during cognitive action control. However, the effect of reward presentation seems subtle and needs further investigation to determine more precisely its relevance. For instance, future studies could rely on recordings of both STN-LFPs and cortical EEG while performing a motivated conflict task. Investigating functional connectivity between the STN and cortical areas such as the orbitofrontal cortex and fronto-parietal networks involved in motivation and cognitive action control would certainly provide very useful information to disentangle the mechanisms by which the STN uses reward information in its computations during conflict resolution.

## Acknowledgements

The authors would like to thank all the patients who took part in this study.

## Abbreviations

DBS: deep brain stimulation
ITPC: intertrial phase clustering
LFP: Local Field Potential
PD: Parkinson’s disease
STN: subthalamic nucleus
UPDRS: Unified Parkinson’s Disease Rating Scale
S&E: Schwab & England
H&Y: Hoehn & Yahr
LEDD: levodopa-equivalent daily dose
MDRS: Mattis Dementia Rating Scale

